# Leveraging multivariate information for community detection in functional brain networks

**DOI:** 10.1101/2024.07.22.604675

**Authors:** Maria Grazia Puxeddu, Maria Pope, Thomas F. Varley, Joshua Faskowitz, Olaf Sporns

## Abstract

Embedded in neuroscience is the concept that brain functioning is underpinned by specialized systems whose integration enables cognition and behavior. Modeling the brain as a network of interconnected brain regions, allowed us to capitalize on network science tools and identify these segregated systems (modules, or communities) by optimizing the weights of pairwise connections within them. However, just knowing how strongly two brain areas are connected does not paint the whole picture. Brain dynamics is also engendered by interactions involving more areas at the same time, namely, higher-order interactions. In this paper, we propose a community detection algorithm that accounts for higher-order interactions and finds modules of brain regions whose brain activity is maximally redundant. Compared to modules identified with methods based on bivariate interactions, our redundancy-dominated modules are more symmetrical between the hemispheres, they overlap with canonical systems at the level of the sensory cortex, but describe a new organization of the transmodal cortex. By detecting redundant modules across spatial scales, we identified a sweet spot of maximum balance between segregation and integration of information as that scale where redundancy within modules and synergy between modules peaked. Moreover, we defined a local index that distinguishes brain regions in segregators and integrators based on how much they participate in the redundancy of their modules versus the redundancy of the whole system. Finally, we applied the algorithm to a lifespan dataset and tracked how redundant subsystems change across time. The results of this paper serve as educated guesses on how the brain organizes itself into modules accounting for higher-order interactions of its fundamental units, and pave the way for further investigation that could link them to cognition, behavior, and disease.

## INTRODUCTION

Emergent properties of complex systems arise from a balance between segregation and integration of the system’s fundamental units [1]. A prominent example of such a system is the brain, which displays ongoing transitions between segregated and integrated activity [2–8]. On the one hand, functional specialization pushes the brain to segregate functionally related groups of neurons, neural populations, or brain areas. On the other hand, the integration of these systems promotes global communication required for coherent perception and behavior. Thus, identifying the principles able to model and recapitulate this interplay is a key goal of computational cognitive neuroscience.

The balance between local segregation and global integration can be viewed through the lens of network science. Under the network conceptualization, the brain can be modeled as an ensemble of distributed brain regions (nodes) linked by anatomical connections (anatomical networks) or dynamic interactions (functional networks) [9, 10]. Segregation in functional brain networks has been studied primarily by observing how nodes are organized into *modules* [11, 12]. Network modules, also referred to as communities or clusters, are groups of nodes that form strong (or dense) connections to one another and weak (or sparse) connections to other nodes in the network. Strongly connected brain modules map out functional systems that are often invoked as building blocks of cognition and behavior [13]. Complementing brain modules are network hubs – nodes highly interconnected with the whole network – which enable information transmission between modules and thus functional integration [14, 15].

The identification of modules, commonly called community detection, is usually conducted using the so-called functional connectivity matrix—the matrix formed by measuring covariance between pairs of neural elements. While valid, this approach is also limited in scope. By design, most community detection techniques search for groupings of elements according to pairwise similarities, without taking into account that more than two neural units can (and do) engage with each other, enabling higher-order interactions [16–18] An increasing body of literature centers on these higher-order interactions as key features of complex systems, including the brain [19–23]. This surge in interest demands new tools able to reveal brain organizational principles by leveraging this augmented space of possible interactions.

In the current paper, we tackle this issue by introducing a new framework to model and investigate how segregation emerges from higher-order interactions of brain regions. We build our work exploiting recent advances in information theory, which provides the mathematics to describe polyadic dependencies in multivariate systems [24]. Given a set of brain regions and their activity portrayed through time series, information theory can help us discern two types of interactions: redundant and synergistic [25]. While *redundant information* captures how much information is copied across the elements (i.e., brain regions), *synergistic information* is the information that is accessible only by considering the state of all the elements as a whole [26]. We postulate that the way the brain organizes itself into segregated subsystems can be captured by higher-order interactions within subsets of brain regions whose activity is maximally redundant, jointly sharing the same information. How information is integrated over the whole brain system can then be estimated by quantifying the level of synergy between elements of different subsets.

For this purpose, we developed a community detection algorithm that finds maximally redundant modules at multiple spatial scales. We assessed how they relate to canonical functional systems and to the modules identified with conventional methods that only consider pairwise interactions. We also quantified the extent to which the information carried by the subsets is integrated over the entire system through synergistic interactions. Moreover, after describing where these subsets are located in the cortex, we also defined a new local index of segregation and integration, based on subset interactions. Finally, we applied our algorithm to a human lifespan fMRI dataset to observe how this higher-order redundancy-dominated modular structure evolves from youth to senescence. Collectively, our work represents a methodological advancement towards a more comprehensive characterization of organizational principles in multivariate systems and its application provides new insights into how the brain organizes itself into functionally segregated subsystems.

## RESULTS

Numerous studies investigated the modular structure of functional brain networks. One challenge is to move beyond the pairwise representation to reveal a more realistic organization of the brain’s dynamics that includes multivariate interactions among multiple brain areas. Here, we address this issue by introducing an algorithm that groups together brain areas that share a significant amount of identical information in their neural activity, that is, we group together maximally redundant brain areas.

Throughout the analyses we used functional connectivity (FC) data derived from resting-state fMRI recordings of three different datasets: the Human Connectome Project (HCP) [27], the Microstructure-Informed Connectomics (MICA-MICs) data [28], and the Nathan Kline Institute (NKI) lifespan data [29]. As in prior work [20], we relyed on Gaussian assumptions to estimate multivariate information theoretic measures from covariance matrices encoding FC. To this end, in each dataset the cerebral cortex was parcellated in 200 nodes [30] and covariance matrices were derived from time series representing BOLD signals. For more information about the datasets and how we retrieved relevant signals, see Methods.

### Community detection via Total Correlation maximization

The most popular methods for community detection group together nodes based on the density and strength of their pairwise interactions. Among these, modularity maximization [31], implemented with the Louvain algorithm [32], is the most widely used in neuroscience applications. It is based on a quality function called ‘modularity’ (Q) that measures the goodness of a partition by counting how many connections fit within a given set of modules compared to chance, i.e., what we would observe in a null network model partitioned in the same way. The Q heuristic can be optimized outright to discover modular community structure and the optimization can be tuned to find modules at different spatial scales, from coarser to finer (see also methods “Multiscale modularity maximization”). While modularity maximization has been proven useful in linking brain network topology to function and behavior, the communities that it finds do not account for higher-order interactions among subsets of nodes and therefore might miss important multivariate aspects of brain organization. Here, we overcame this issue by introducing a community detection algorithm that leverages higher-order interactions.

Information theory proposes the total correlation (TC) as a proxy for redundancy [1, 33–35]. Given a multivariate system, TC is low when every element is independent and high when the joint state of the elements has low entropy, given the individual entropies, i.e., the system is dominated by redundant interactions. For a full mathematical description of TC see Equations 7 and 13 in Methods. We hypothesized that modules are made of maximally redundant brain areas, that is, sets of nodes that share a large amount of information. Thus, we introduced the total correlation score (TC_score_), a new quality function that, given a partition of the brain into subsets, estimates the redundancy of the modules compared to chance:

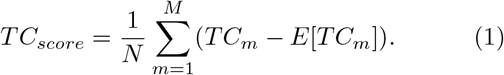

with *N* being the number of nodes, *M* the number of modules, *TC*_*m*_ the total correlation within module *m*, and *E*[*TC*_*m*_] the mean level of integration for randomly selected subsets of the same size of module *m*. Thus, TC_score_ is higher the more the TC within modules exceeds the average TC of equal size subsets. High TC_score_ values entail a good partition, whereas for particularly poor partitions, or very weakly integrated systems, this quality function can even be negative.

Analogously to Q, TC_score_ can be used to assess the goodness of a partition, or it can be maximized to infer an optimal modular structure. Here, we implemented an optimization algorithm based on simulated annealing, for which we report the pseudocode and a schematic in Table I and Figure 1. The first step of the algorithm consists of computing the TSE complexity curve [1]. This provides the maximum and mean levels of integration for any subset size for the network that we want to partition (all possible E[TC_m_] in Eq.1, and see Figure 1A). Starting from a random (seed) partition then, we can compute the TC of the subsets identified by this partition and compute TC_score_. At this point, the algorithm randomly switches node assignments (without changing the number of modules) and uses simulated annealing to search in the space of solutions for a partition that maximizes TC_score_.

**TABLE I.**
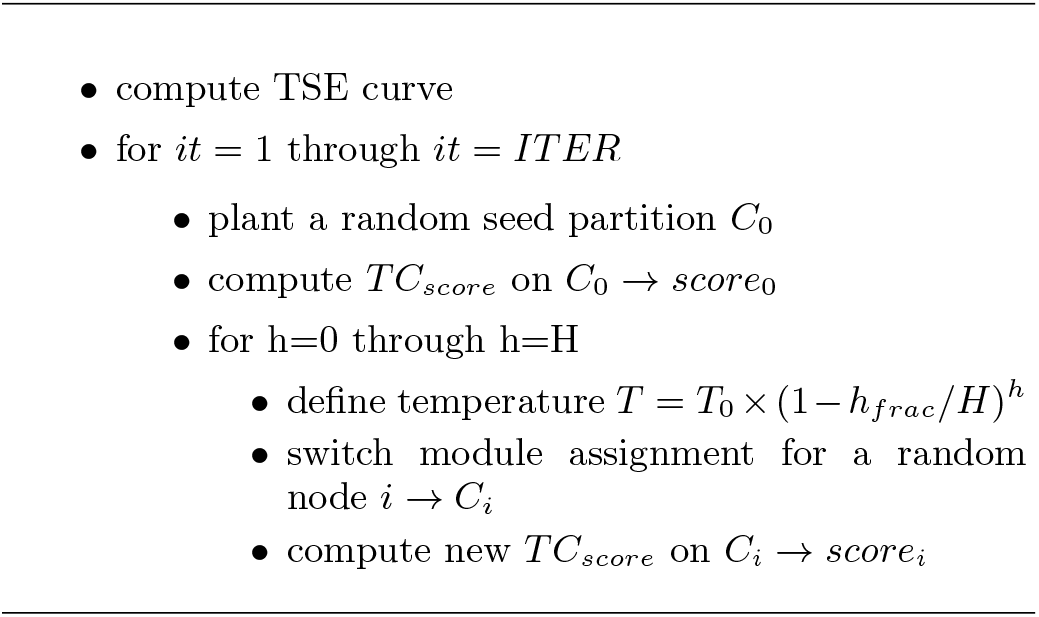
Pseudocode of our algorithm for community detection via total correlation optimization. We set the parameters as follows: H=100000 (number of iterations in the annealing process); ITER=100 (number of times we perform the optimization through annealing); *h*_*frac*_ = 10; *T*_0_ = 1.

**FIG. 1.**
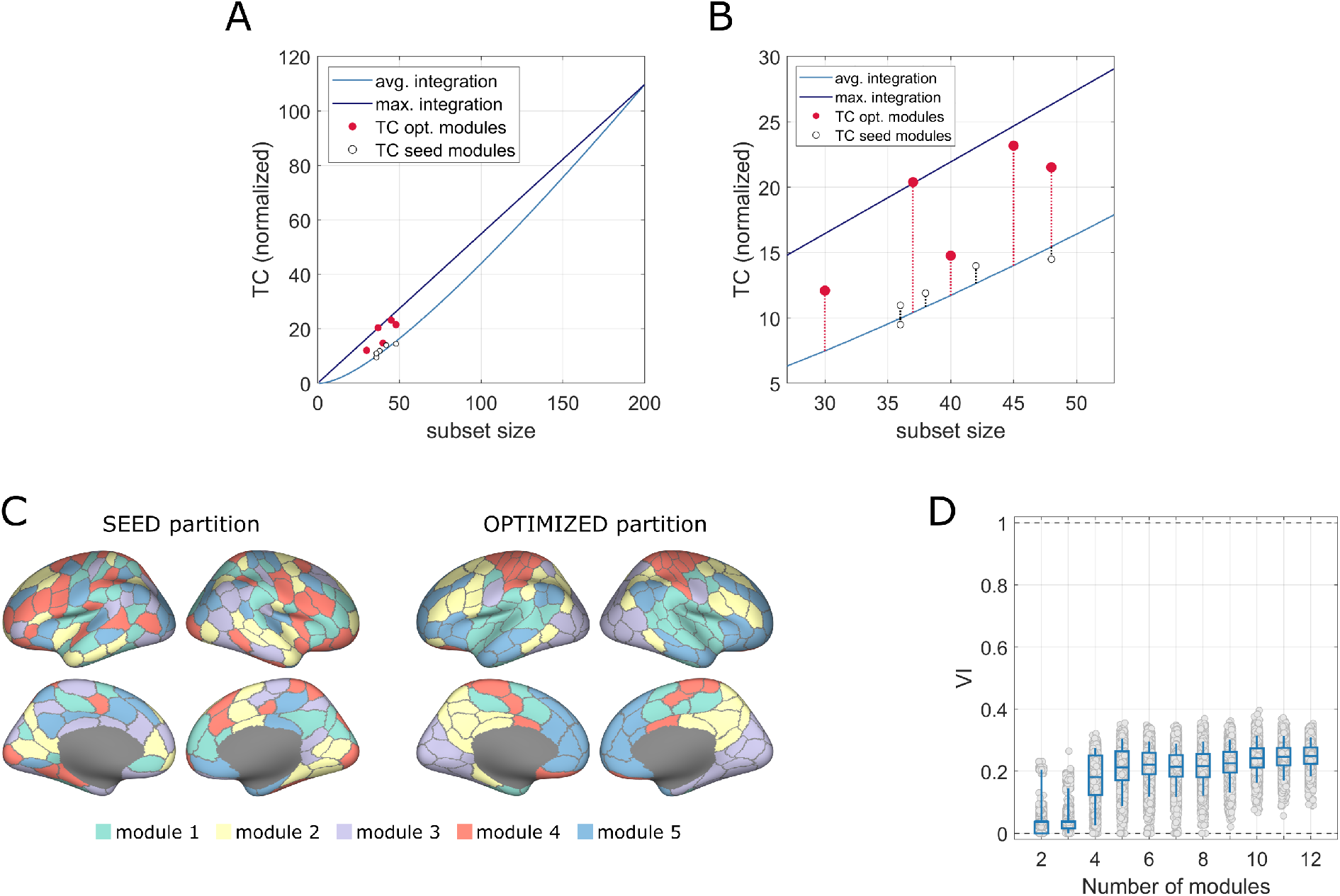
Schematic of the algorithm. **A**. TSE curves for the HCP covariance matrix. For any subset size, they provide an estimate of the average (blue) and maximum (purple) levels of integration of the nodes in the subsets in terms of total correlation. By partitioning the system into modules, we can compute the TC of the modules as the TC of the nodes constituting each module. Then, we can map them in the same graph to see where these values are located with respect to the average level of integration for subsets of the same size. The white dots are the TC of the modules of a random partition, whereas the red dots are the TC of the modules of the optimized partition. **B**. A zoomed in version of panel A, which clarifies that in the optimized partition the modules have a much higher level of integration. **C**. An example of a random seed partition and the partition obtained after the optimization, projected on the cortex. **D**. How similar the partitions obtained from different runs of the optimization are, measured at each spatial scale by using variation of information (VI) as the similarity metric.

In Figure 1C we report an example of what we obtained by running the algorithm on the HCP data. We projected on the cortex an initial random partition and the partition that optimizes TC_score_. Analogously, we display in Figure 1B the TC of the initial random modules and the TC of the optimized modules, which is much higher compared to the average TC of subsets of equivalent size. Finally, we ran the algorithm 100 times, varying the number of modules from 2 to 12. We computed the similarity between the 100 optimized partitions at each resolution in terms of variation of information (VI) [36]. Low VI values suggest that the algorithm delivered partitions that were highly similar and consistent across multiple attempts (Figure 1D).

### Relation between TCscore and Q

How do the two heuristics, TC_score_ and Q, relate to each other? To answer this question, we applied multiresolution consensus clustering (MRCC) [37] to the HCP FC covariance matrix and we computed the TC_score_ on the resulting partitions. MRCC uses the Louvain algorithm to optimize Q at different spatial scales, thus providing partitions made of finer and coarser modules. By running MRCC, we obtained 990 partitions made of a number of modules within the range [2, 50] (see also Methods “Multiscale modularity maximization”). Each one of these partitions is associated with a Q-value that resulted from the optimization, which we compared to the TC_score_ values computed on the same partitions. TC_score_ and Q are significantly and strongly correlated (*r* = 0.91; *pval <* 10 15; Figure 2A). However, if we divide the ensemble of data points into groups where partitions have the same number of modules, TC_score_ and Q result anti-correlated (Figure 2B). This means that TC_score_ and Q are linked by a Simpson’s paradox: overall the two variables are positively correlated, but this correlation is reversed when the ensemble is divided into groups. Specifically, both TC_score_ and Q values increase for coarser partitions (i.e., partitions made of fewer modules), but if we consider partitions with equal levels of granularity the partition with higher TC_score_ is the one with lower Q and vice versa.

**FIG. 2.**
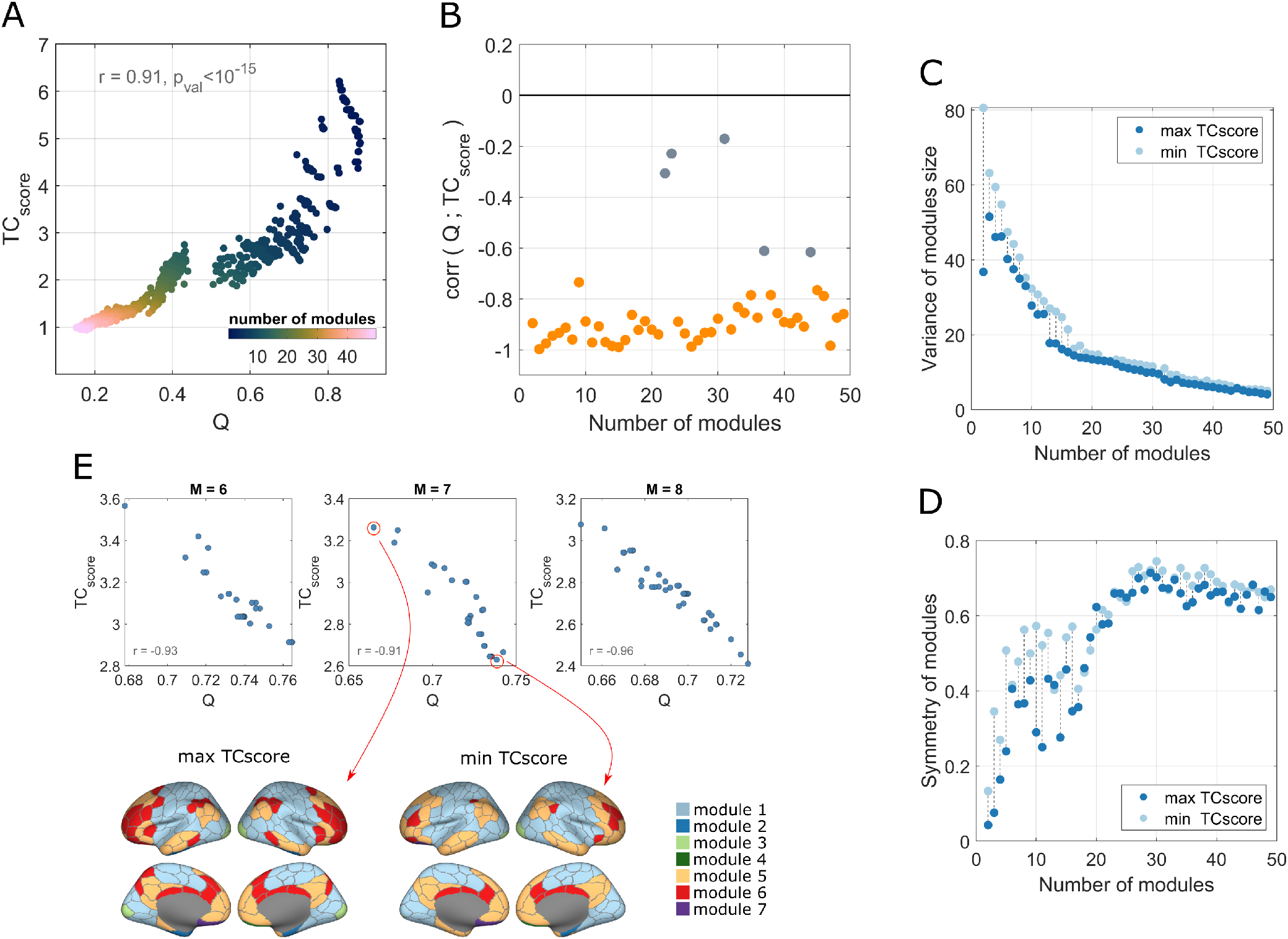
Relation between TC_score_ and Q. **A**. Trend of TC_score_ relative to Q. Each point corresponds to a partition obtained optimizing Q on the HCP FC network and on which we computed TC_score_ downstream. Colors code the spatial resolution of the partitions indicating the number of clusters. **B**. For each group of partitions with an equal number of modules, we computed the correlation between TC_score_ and Q and reported here the correlation coefficients. Points colored in orange identify statistically significant correlations. **C-D**. Values of variance of the modules size and symmetry between hemispheres at each spatial resolution for the two partitions with highest and lowest TC_score_. **E**. Example of how the positive correlation observed in panel A is reversed if we consider groups of partitions with equal resolution (or equal number of modules). In this case, we report the reversed correlation for groups of partitions with 6, 7, and 8 modules. We also report a projection on the cortex surface of the modular structure of the 7-module partitions with the highest and lowest TC_score_.

We explored what might cause the anticorrelation within partition groups. For each set of partitions with an equal number of modules we considered the two partitions with the highest and lowest TC_score_ (Figure 2E) and computed the variance of the modules size and the symmetry of the modules between the hemispheres. The variance of modules size was computed as the standard deviation of the size of the modules (i.e., how many nodes belong to one module) so that higher values mean that the sizes of modules are heterogeneous within the partition. To compute the symmetry between hemispheres, we used a measure of “unbalance”, which for each module counts the difference between the number of nodes on the left hemisphere and those on the right hemisphere, normalized with the total number of nodes. The higher this measure, the lower the symmetry of the module with respect to the hemispheres. The results, reported in Figure 2C-D, show that the partitions with the lowest TC_score_ (higher Q) consistently present modules with greater variance and a lower hemispheric symmetry, compared to the partitions with the highest TC_score_ (lowest Q).

Altogether, these findings suggest that our newly introduced TC_score_ fairly recapitulates Q in some aspects. However, TC_score_ rewards modular structures that are more symmetric between cortical hemispheres and composed of modules of similar size.

### Relation with the canonical systems

In the previous section, we used TC_score_ as a quality function to assess the goodness of partitions found with modularity maximization to see how well our metric recapitulates conventionally detected modular structures. Here, we use our algorithm to maximize TC_score_ on FC networks. One question when inferring a new modular structure in the human brain is how it relates to resting state functional systems (RSFS). Throughout the neuroimaging literature, different labels have been applied to RSFC communities, nonetheless, the patterns of organization have been largely consistent [38–40]. However, all these studies have always overlooked higher-order interactions among brain regions, at least in terms of redundancy and synergy. In this section, we report how the modules derived from TCscore maximization, relate to previously reported RSFS, and specifically to well-established canonical systems [39].

We ran our algorithm initializing it with partition seeds characterized by a different number of communities (M=[2, 12]), so that also the number of communities of the optimized partitions varied across spatial scales. For each spatial resolution we ran the algorithm 100 times and, for each one of these iterations, we computed the similarity between the detected partition and the partition corresponding to the canonical systems. As an index of similarity, we used the Adjusted Mutual Information (AMI) [41], a measure that theoretically accounts for differences in the number of modules between two partitions. The AMI is bounded in the range [0, 1], with higher values indicating higer similarity (AMI=1 means identical partitions). As reported in Figure 3A, at each resolution the optimized partitions present some level of overlap with the canonical systems (the AMI is never 0), but at the same time, they are never identical (the AMI is never 1). As expected, the maximum similarity is obtained when M=7, that is the number of the canonical systems.

**FIG. 3.**
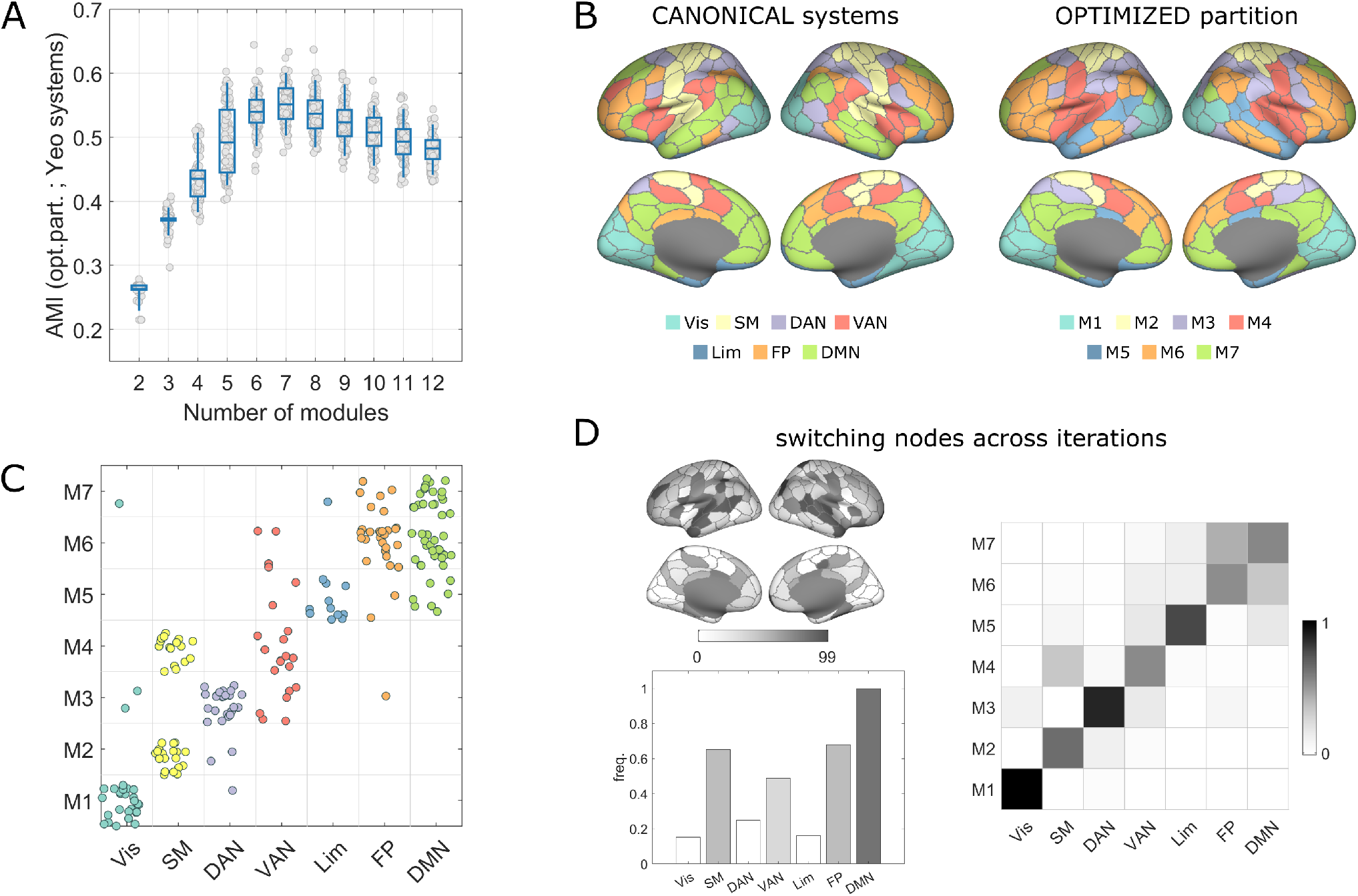
Relation with the canonical RSFS. **A**. Similarity between the partitions inferred by maximizing TC_score_, and the canonical systems in terms of Adjusted Mutual Information. The boxplots summarize the statistics of the 100 iterations of the algorithm at each spatial scale. **B**. Projection on the cortex surface of the canonical systems (left) and the partition obtained optimizing TC_score_ (right). For the representation of the latter, we report the centroid partition (the partition most similar to all the others within the 100 iterations of the optimization algorithm) with 7 modules. **C**. In this panel, each dot is a node and we show to which module/system it is assigned in the two partitions shown in panel B. The colors follow the RSFS classification. **D**. We further measure the overlap considering the whole set of 100 partitions obtained in the 100 iterations of the optimization. On the left, we report the frequency with which nodes change module assignments between the canonical systems and each one of the 100 partitions. We report this frequency at the node-level on the cortex and at the system-level through the bar plot. Darker greys depict nodes/systems whose assignment varies the most between the canonical systems and the optimized partitions. On the right, we report the same visualization of panel C, but considering the 100 iterations; darker squares represent a larger overlap between modules across all the iterations.

Next, we investigated the relation of our partitions to canonical systems in more detail. We focused on TCoptimized partitions with 7 modules. A first visual comparison can be drawn from Figure 3B, where we represented a projection on the cortex of the two partitions. We noticed that modules largely overlap, with some differences. We further mapped *how* they overlap by module/system in Figure 3C, demonstrating that the greatest correspondence between module assignment is found for the visual system, the dorsal attention network (DAN), and the limbic system, which are well captured by modules 1, 3, and 5 of our optimized partition. The somatomotor system (SM) splits into modules 2 and 4, and part of it is co-assigned to the ventral attention system (VAN). Interestingly, recent studies highlighted high levels of co-activations between the two [42– The default mode network (DMN) and frontoparietal networks (FP) instead are mixed into modules 6 and 7, suggesting that association areas are redistributed when considering higher-order interactions in the community detection process. These results are consistent among the 100 iterations of the algorithm (Figure 3D), and are neatly replicated in the MICA dataset (Figure S1, A-D).

Altogether these results suggest that partitions inferred by considering higher-order interactions via TC_score_ maximization partially preserve the organization recapitulated by the familiar canonical systems. However, they also suggest a different organization for regions comprising higher order association systems.

### Redundancy dominated modules across spatial scales

In this section, we provide a more thorough description of the modular structure inferred by maximizing TC_score_, analyzing how it balances segregation and integration of information at different spatial resolutions. For this purpose, we used the same set of optimized partitions of the previous analysis. This set comprises 100 instances for each spatial scale (identified by the number of clusters, M=[2, 12]), derived from the HCP FC covariance matrices.

First, we investigated how TC_score_ varies across spatial scales. It showed a decreasing trend with respect to M (Figure 4A), with a peak in correspondence of partitions made of 3 modules. This means that, as the partitions become finer and modules smaller, modules are comprised of nodes carrying less redundant information, i.e., they become less segregated.

**FIG. 4.**
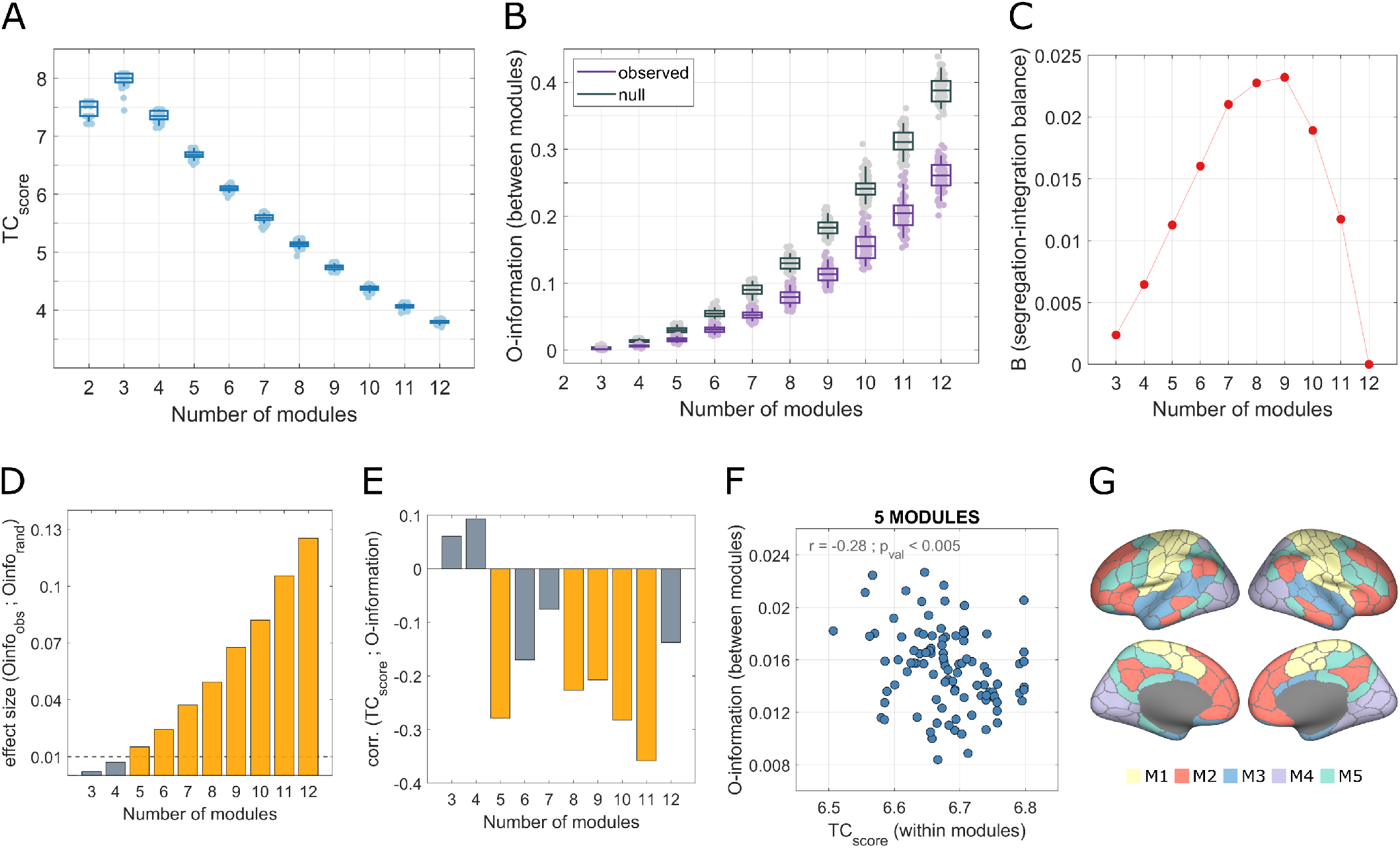
**A**. Trend of TC_score_ versus the number of clusters. The statistics to display the boxplots were carried out considering 100 iterations of the optimization algorithm for each spatial scale. **B**. Trend of O-information versus number of clusters in the optimized partitions (purple) and in randomized nodes (gray). **C**. Trend of B, the index that recapitulates the balance between redundancy within modules and synergy between modules (segregation-integration balance in a higher order space), across spatial resolutions **D**. Median difference effect size of the observations from the two distributions of empirical and null model-derived O-information. **E**. Correlation coefficients obtained by computing the Pearson correlation between TC_score_ and O-information among the 100 iterations at each spatial scale. Statistically significant correlations have been highlighted in yellow. **F**. Example of such correlation for M=5. **G**. Projection on the cortex of the 5-module partition with highest TC_score_ and lowest O-information.

Then, we focused on how the information carried by single modules is integrated throughout the system at different spatial scales. We hypothesized that an informationally optimal community structure is one where modules carry distinct information that becomes accessible when observed together. This phenomenon is well captured by the notion of synergy, which we computationally quantify through the organizational information, or O-information [24]. In a multivariate system, the lower the O-information the more the information carried by a set of variables is synergistic (for details see Methods “Synergistic and redundant information in multivariate systems”). To have an estimate of how synergistic infor-mation manifests across modules, we randomly sampled nodes from different modules, for a total of 1000 times for each spatial scale, and computed the O-information on the obtained subsets. For instance, in partitions made of 7 modules, we sampled 7 nodes, each belonging to a different module, and computed the O-information in such subset. We carried out the same analysis on a null model, where we sampled the same number of nodes but at random, without considering the modular structure. Whereas TC_score_ is defined relative to a null model (the average level of integration in the TSE curve), the Oinformation lacks a similar comparison in its formutation, thus necessitating this extra null model comparison. We observed that the information carried by different modules is more synergistic in coarser partitions (lower O-information), however, when the partitions become finer, the O-information deviates more from the null model (Figure 4B). We quantified this deviation by computing the effect size (ES) as the difference between the means of the observed and the null O-information (Figure 4D). This metric captures that in finer partitions the amount of synergistic information retrievable in a subset of nodes is more closely linked to the community structure – nodes from different modules actually carry distinct information – whereas at a coarser scale we might obtain greater synergy just because it is computed on a smaller number of elements.

Next, we investigated the relationship between TC_score_ and O-information for specific spatial scales. For any given resolution, we computed the correlation between the two variables across the 100 iterations (Figure 4E,G). For most of the scales, TC_score_ and O-information were significantly anti-correlated: the partitions that have maximally redundant modules are also those where the synergy between modules is the greatest. This balance of redundancy within and synergy between modules can be view as an expression of the co-existence of segregation and integration in a higher-order space.

Once we observed the trends of redundancy and synergy across scales, we asked: is there an optimal scale where redundancy within and synergy between modules are maximally balanced? To answer this question, we defined the segregation-integration balance index (B) as follows:

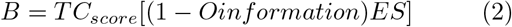

Ideally, we want the B to be as high as possible: when the three factors are high, we have maximum redundancy within and maximum synergy between modules. In other words, each module is made of nodes that share a large amount of information (high TC_score_) and carry different information to the network (high (1Oinformation) × ES). Tracking this index along spatial resolutions (Figure 4C), resulted in a bell shape with a peak in M=9 and highest values in the range M=[7, 9]. This is a direct consequence of what observed in Figure 4A,B. The redundancy within modules is higher in coarser partitions. The synergy between modules is also higher in coarser partitions, however, it deviates more from what we would observe in a null model (measured by ES) as the partitions become finer. Thus, it is intuitive that a combination of redundancy and synergy, that captures the higher-order balance between segregation and integration, can be observed at intermediate scales, when partitions are neither too coarse nor too fine.

Altogether, these results suggest that redundancy and synergy of brain activity within and between modules, and their interplay, vary across spatial resolution. An optimal balance has been found in partitions composed of 7 to 9 modules. All the findings are replicated in the MICA dataset (Figure S1, E-H).

### Local properties of the redundancy dominated modular structure

Having defined globally in which way the new modular structure balances redundant and synergistic information, we now want to characterize its local properties. Specifically, we want to investigate how much single brain regions participate in the redundant integration of the module to which they belong, relative to how much they participate in the redundant integration of the whole network. This concept is analogous to what is commonly known in network science as *participation coefficient* [45], but here we include the consideration of higher-order interactions. While computing nodal measures in bivariate networks is straightforward, it becomes more difficult when considering multivariate measures such as TC, which by definition, cannot be computed on single nodes.

To address this issue, we defined the Relative Integration Coefficient (RIC), which can be computed for each node *i*, and that is formulated as follows:

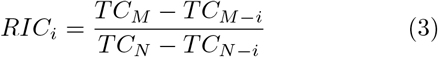

Where *TC*_*M*_ is the *TC* computed on the module *M* to which node *i* belongs, *TC*_*M*−*i*_ is the *TC* of the same module when node *i* is removed, *TC*_*N*_ is the *TC* computed on the whole network, and *TC*_*N*−*i*_ is the *TC* computed on the whole network after removing node *i*. The idea is that, even if we cannot compute TC on single nodes, we can look at how much the TC increases (or decreases) when we remove node i from the modules and the network, in order to quantify how much it participates in the redundancy of its module compared to the redundancy of the whole system. High RIC-values indicate that the node contributes to the redundancy of the module more than to the redundancy of the network (if we remove it, the TC of the module becomes low, hence the numerator is high). We call nodes with high RIC *segregator* nodes, as their role is to share information predominantly with nodes of the same module. This is particularly true when RIC>1. On the contrary, low values of RIC indicate that the node highly contributes to the redundancy of the whole network (if we remove it the TC of the network becomes high, hence the denominator is low). We call these nodes *integrators*, as their role is to share information with nodes of the entire network.

In Figure 5A we displayed the RIC distribution computed on the centroid partition of the 100 instantiations with 7 modules. It shows a left tail of integrator nodes (low RIC-values), and a right tail of segregator nodes (RIC>1). In Figure 5B we localized segregator and integrator nodes on the cortex and with respect to the canonical systems. Segregators reside mainly in the visual and somatomotor systems, i.e., in the sensory areas. Conversely, integrators can be found mostly in the DMN, FP, and VAN, i.e., in the transmodal and association cortex. This is plausible given their role of integrating information from different areas to perform cognitive functions. This local mapping is consistent across the 100 iterations of the algorithm and spatial scales (Figure 5C,D). Moreover, a replication on the MICA dataset shows coherent results (Figure S1,I-K).

**FIG. 5.**
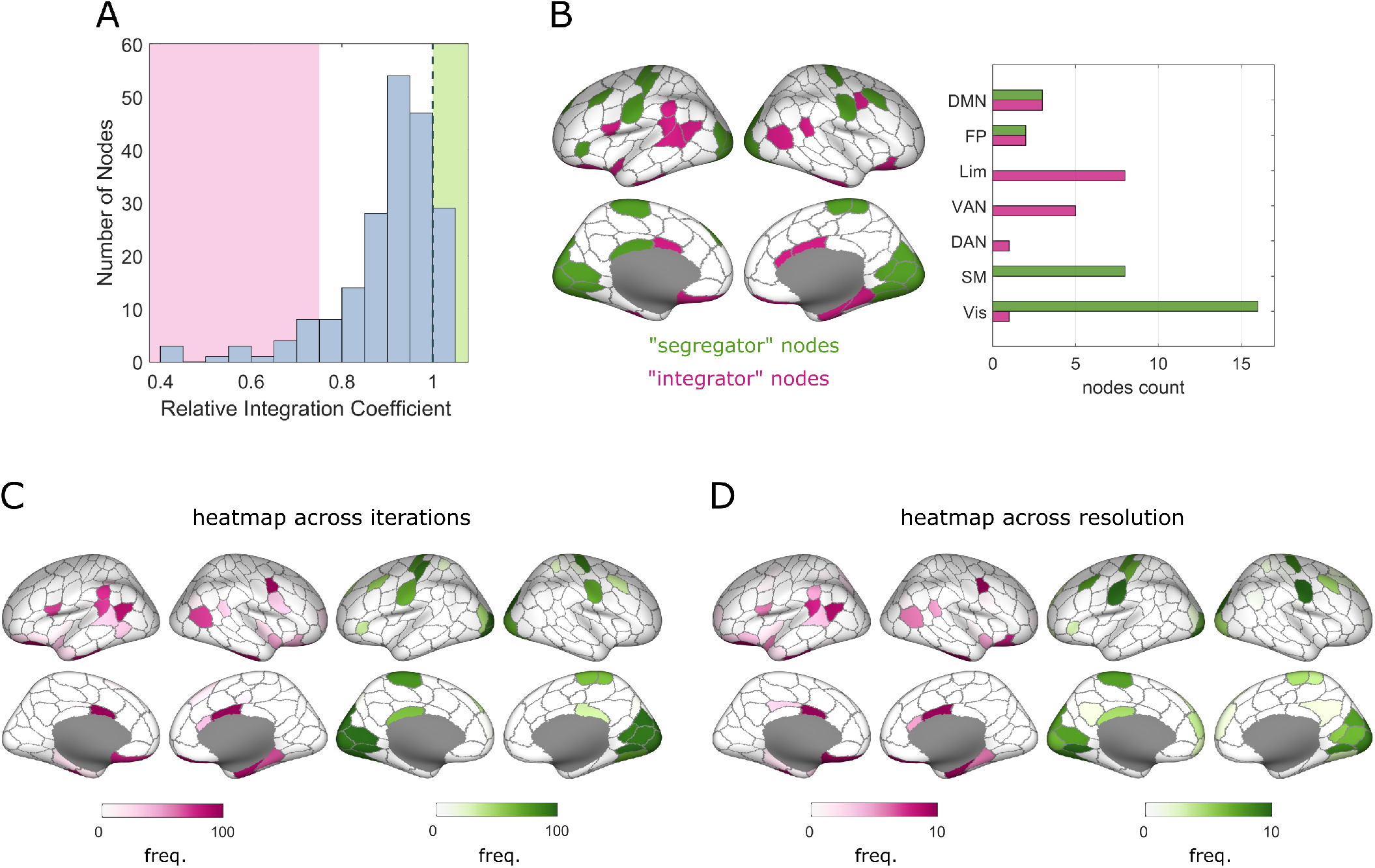
Relative Integration Coefficient. **A**. Histogram of relative integration coefficient computed for each node. The area where RIC is greater than 1, denoting segregator nodes, has been highlighted in green, whereas the area containing 10% of the lowest RIC values, denoting integrator nodes, has been colored in pink. **B**.Projection on the cortex of the nodes identified as segregator (green) and integrator (pink) in the centroid partition with 7 modules and also count of them within the canonical systems (on the right). **C-D**. Projection on the cortex of the frequency with which nodes have been counted as segregator or integrator among the 100 partitions with 7 modules (panel C) and the centroids partitions across spatial scales (panel D). The intensity of the color is proportional to that frequency.

Ultimately, with this local analysis, we could assign a role to single nodes, derived from their participation in higher-order interactions relative to the modular structure. We were able to identify brain areas more devoted to sharing information with similar nodes (i.e., nodes belonging to the same community) as well as brain areas whose primary purpose is integrating information across the whole network.

### Redundancy dominated modular structure across the human lifespan

Lastly, we wanted to test the ability of TC-based community detection to capture variations in functional connectivity organization that could naturally arise when considering clinical cohorts, or large-scale multi-subject datasets. To this purpose, we applied our new methodology to a lifespan dataset, as the progressive modifications exhibited during development, adulthood, and senescence, are one of the most striking and well-documented examples of functional connectivity re-organization [46–48].

We leveraged the NKI dataset [29], comprised of 917 subjects with ages heterogeneously distributed between 6 to 85 years (Figure S2A). For each one of the subjects, we performed 100 iterations of the TC_score_ optimization at different spatial resolutions, obtaining partitions with a number of modules between 2 and 10. Here we reported the results of the analysis for M=9 (for each subject we considered the centroid partition after iterating the algorithm 100 times), as we showed that it is the resolution where segregation and integration of information are optimally balanced. However, the analyses have been carried out at every spatial scale and the results, which were consistent with those presented in the main text, can be found in Supplementary Information.

First, we inspected how redundancy within modules and synergy among them vary across the lifespan (Figure 6A,B). We found that TC_score_ significantly decreases with age 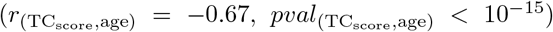, whereas O-information between modules significantly increases (*r*_(Oinformation,age)_ = 0.40, *pval*_(Oinformation,age)_ < 10^−15^). This result, (confirmed at different spatial scales, Figure S4) suggests that aging is associated with a decrease of segregation of the network, a phenomenon largely observed in previous studies [12]. Moreover, aging is associated with less stability of the algorithm (Figure S3), which might indicate that it is more difficult to retrieve maximally redundant modules in older adults.

**FIG. 6.**
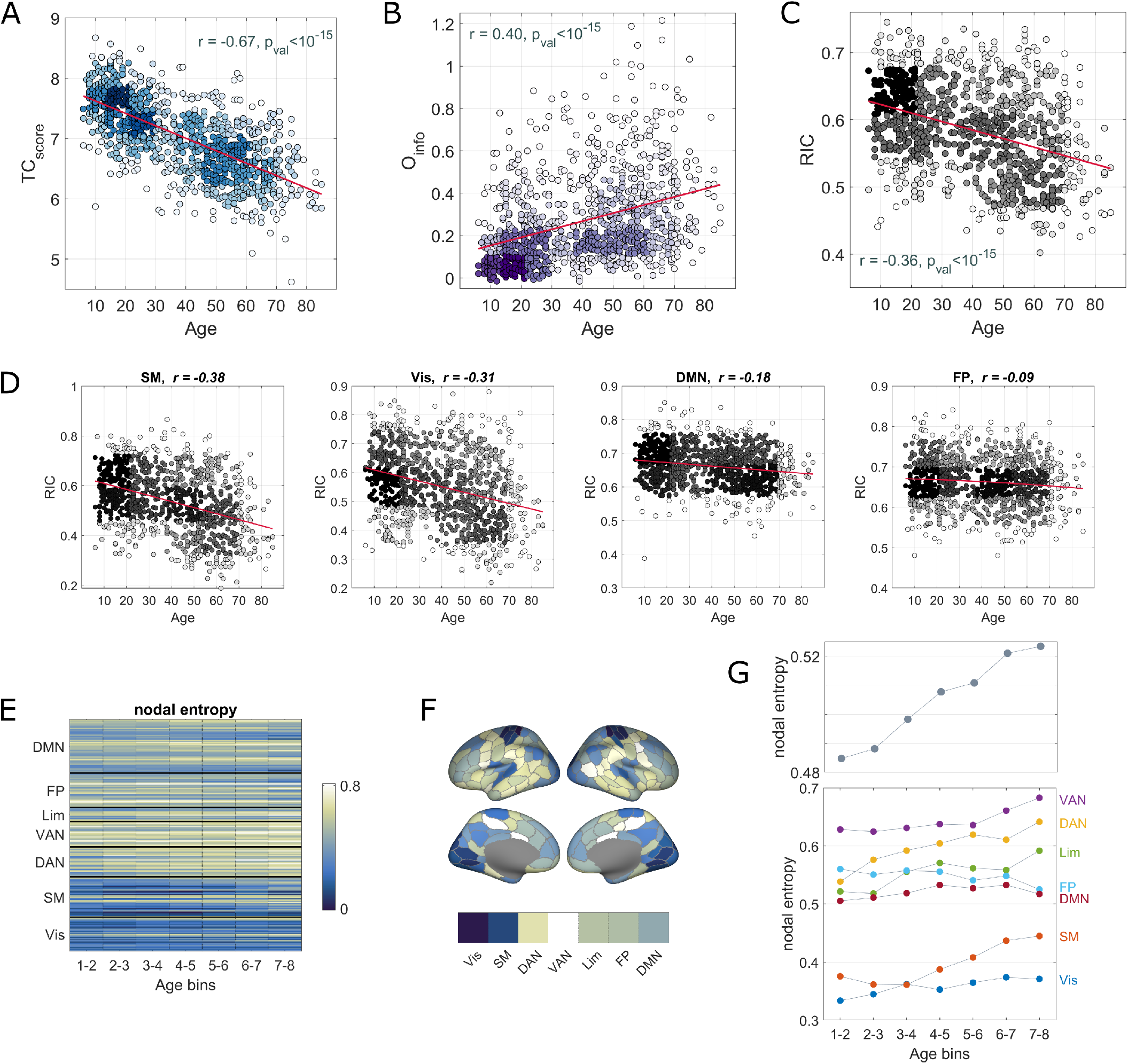
Application to the NKI dataset. **A**. Values of the TC_score_ for the partitions inferred at the single-subject level, versus age. **B**.Values of O-information between modules computed on single-subject partitions versus age. **C**. RIC values averaged across nodes for each subject versus age. **D**. RIC values averaged within the canonical systems, for each subject, versus age. Throughout panels A-D, because the population is not uniformly distributed across age, we represented values in age bins with higher population density with darker colors. **E**. With the nodal entropy we depict how much single nodes change module assignment across the lifespan. Yellow colors indicate high entropy, i.e., nodes that are more likely to change module, and vice versa colors toward blue indicate nodes whose module assignment is stable. **F**. Projection on the cortex surface of the nodal entropy, averaged across age-groups. **G**. Values of nodal entropy averaged within the canonical systems and reported for each age group. In the upper graph the value of nodal entropy, averaged across nodes, is reported.

Next, we computed the RIC-values of each node, thus mapping age-related varions of the extent to which a node participates to the redundancy of its module. We found that the average RIC values decrease with age (Figure 6C), meaning that later in life nodes participate more to the integration of the network, rather than individual modules. Unpacking the RIC’s trend for each canonical system (Figure 6D), we observed that nodes in the somatomotor and visual systems, previously classified as segregator nodes, have the steepest decline of RIC, which means that they drive the progressive integration of the network. On the other hand, nodes from the DMN and FP networks, classified as integrator nodes, show a modest decline of RIC. Nodes from other canonical systems, not reported in the figure, presented an intermediate pat-tern.

A final question is how much and in what way the modular structure changes across the lifespan. To answer this question, we divided the dataset into 8 age groups, each spanning 10 years. Analogously to what has been done in [49], we used a bootstrap strategy where for 500 iterations we sampled subjects in each group, to then identify which nodes change allegiance to modules across consecutive groups. We quantified this change by re-aligning the labels of the partitions with respect to those of the first age-group and then counting the mismatched labels across age-groups. We called the average of this measure across the 500 iterations *nodal entropy* (the higher it is, the more that node changes module assignment). We observed that the nodes belonging to the visual and somatomotor systems are the most stable across the lifespan (Figure 6E,F). We also averaged this measure within the canonical systems (Figure 6G), finding that in the late lifespan, the reconfiguration of the modular structure is greater than at the beginning (i.e., the entropy increases). Most of the systems reconfigure more with age, except for the DMN and the FP networks, which keep the most stable levels of nodal entropy.

Overall, these results suggest that our newly introduced algorithm is able to track changes in functional organization. The sensory areas overlap with the most stable modules. However, the changes that they present are aimed at integrating the whole system. The DMN, FP and attention networks are those that reconfigure the most, but they keep stable levels of integration with the whole network from the beginning of the lifespan to the end.

## DISCUSSION

In this paper, we proposed an algorithm that uncovers modular structure in functional brain networks by accounting for higher-order interactions. Our approach leverages the mathematical framework of information theory to find highly redundant subsets of nodes. We provided a new view of brain organization and showed how it contrasts with canonical functional systems and with the modular structure uncovered with traditional methods based on pairwise interactions. The discovery of a Simpson’s paradox relating TC_score_ and modularity suggests that, when higher-order interactions are accounted for, the notion of ‘structure’ in complex systems becomes much richer than is typically observed in purely bivariate systems. This may be a fruitful new perspective, as the Simpson’s paradox typically reflects non-trivial interactions between elements of a system, when viewed relative to certain bounds. We also presented two new indices: a global index that estimates how the redundant modules balance segregation and integration of information across the whole network, and a local index that characterizes node diversity in terms of contribution to network integration. Finally, we delineated the evolution of redundant modular structure across the human lifespan.

### Redundancy dominated systems of the human brain

Decades of neuroimaging research have been devoted to defining functional systems of the human brain. Task-based experiments identified such systems as collections of co-activated brain areas [50, 51]. More recently, the idea that brain function is intrinsic [52], constantly processing information for interpreting and responding to environmental demands, led to a description of functional systems of highly correlated spontaneous activity [38–40] Proof that such systems are grounded in neurophysiology is widespread. Not only are they largely consistent across studies and methodologies, but some of them also match patterns of brain damage observed in clinical cohorts [53]. However, there exist co-fluctuating areas that have not been firmly aligned with independently validated brain systems [12]. Whether they correspond to undefined brain systems or are artifacts induced by network construction is yet to be determined. We argue that accounting for higher-order interactions is an avenue to unravel more complex relationships among brain areas. This can lead to a different perspective on brain organization and elucidate the role of nodes that are more difficult to place in the classical framework.

We compared redundancy dominated modules to canonical resting-state systems. While some level of similarity was expected (canonical systems are dominated by redundant interactions [54]), we highlight how they differ. Our method led to a clear identification of the visual and somatomotor systems, and the dorsal attention network, which are also the systems more consistently detected across reports. However, the somatomotor system is split into two modules. This is consistent with a dorsal and a ventral representation of such system, reported in [38]. In our partition, some brain areas normally associated with the ventral attention system are co-assigned to nodes of the ventral part of the somatomotor area. Interestingly, previous reports showed co-fluctuations among those regions [42–44] This is an example of how beyond pairwise interactions offer a different angle on how co-activated brain regions could form coherent systems, still grounded in neurophysiology. As for the brain areas belonging to the frontoparietal system and DMN, these converge into two redundant modules where they are mixed. Future investigations will clarify the role of these new systems during tasks, cognitive functions, or disease progression.

### A higher-order lens on segregation and integration of brain systems

The modular organization of the human brain supports segregation and integration of information among functional brain systems [3], enabling cognition and behavior.

Recent studies [56] found proof that local within-systems connectivity is critical for motor execution, and between-systems connectivity is crucial for cognitive tasks. Furthermore, brain organization benefits from the coexistence of segregation and integration. While lack of segregation hampers functional specialization and fails to protect the system from the spreading of disease [57], a complete disconnection between brain systems is associated with brain disorders (e.g., Alzheimer’s [58] and Parkinson’s [59] disease, or schizophrenia [60]) and leads to cognitive deficit [61].

In our work, we explored how the balance between segregation and integration is expressed when considering higher-order interactions. Notably, the TSE complexity [1], that inspired our TC_score_, was designed to capture how brain dynamics optimizes segregation and integration of information. However, it did not account for any modular structure, which now we know is central to brain organization. Instead, we hypothesize that the dichotomy of segregation-integration is supported by a mesoscale organization made of subsystems where brain activity is redundant (copied over the system’s elements) and whose interaction with each other leads to a new type of information, which is synergistic, that goes beyond functional and local specialization. Interestingly, we found these two measures to be anti-correlated: the brain is organized such that the more redundant information we find within subsystems, the more synergistic information we find between them. This means that spontaneously and even under a higher-order perspective, brain activity manifests in a manner where segregation and integration are maximized. We further characterized the balance between segregation and integration via an index that tracks how information encoded in brain activity is redundant within subsystems and synergistic between them. We explored how this index is conveyed across spatial resolutions, finding that a division of the neocortex into 9 subsystems optimizes the abovementioned balance, immediately followed by partitions into 8 and 7 subsystems.

### Nodal diversity in redundancy-dominated modular structure

Given a division of the brain into subsystems, individual nodes can behave differently in how much they interact with nodes of the same system versus how much they interact with other nodes assigned to different systems. In this sense we might say that they serve distinct functional roles, especially if they are highly interconnected hubs: they either foster local specialization (provincial hubs), or facilitate inter-module communication (connector hubs). The participation coefficient [45, 62] was introduced to quantify the extent to which they cover one role or the other. Many reports located connector hubs in the association cortex [14, 63, 64] highlighting how they are involved in a wide repertoire of tasks and functions [65, 66].

All these findings build on a dyadic view of the interactions among brain regions. Recent works [67, 68] studied how being a hub affects computation capacity estimated in terms of multivariate information processing, and information theory has also been used to extend the formulation of participation coefficient to the case where multiple sets of partitions can describe the mesoscale organization of a complex system [69]. However, a measure able to capture the role of the nodes in the modular organization, which also accounts for higherorder interactions, is missing. Thus, we introduced the Relative Integration Coefficient. By shifting the concept of participation coefficient to a higher-order space, we identified segregator and integrator nodes based on the amount of information shared with co-assigned and noncoassigned nodes. Integrators mainly reside in the sensory areas, whereas segregators lie in the DMN, ventral attention, and frontoparietal networks. Notably, brain regions whose predominant role is to enhance modular redundancy were also found to be the most synergistic areas in the system, regardless of the modular structure [20, 23]. A similar distinction between sensory and association areas has also been observed in [70], where the study of co-fluctuations between sets of three and four regions saw the sensory areas as maximally coherent, as opposed to association areas. Future works will elucidate how these maximally synergistic, provincial, coherent areas engage during tasks or in different clinical cohorts.

### Brain redundant subsystems have unique trajectories during the lifespan

The application of our algorithm to a lifespan dataset mainly served the purpose of validating its ability to detect changes in modular organization. Nonetheless, understanding how higher-order interactions change across the years, and with them the emerging modular structure, is of crucial importance and here we provided a first glance at it.

Alterations in the organization of brain connectivity have been observed, even in the absence of disease, in anatomical and functional networks, and in their relationship [46–49,71]. Thus, it is safe to assume that multivariate interactions among brain regions are also subject to change over the lifespan. By uncovering redundancy-dominated modules in subjects between 6 and 85 years old, we tested if, and to what extent, our algorithm is able to capture such changes. First, we observed a decrease within-module redundancy, and thus segregation of information, associated with aging. Evidence of this phenomenon is overwhelming in analyses of pairwise connectivity [72–76 but also in those accounting for higher-order interactions [77, 78]. The fact that we could replicate this ensures that our algorithm is able to track changes in modular organization. We then extended our analysis by portraying how the heterogeneity of node processing with respect to the modular organization unravels over the lifespan. We found that segregators became increasingly less so with age, whereas integrators became only mildly more integrated in the system. This suggests that the decreased segregation is majorly driven by sensory areas that share more and more information with other systems. This might align with a reduction in the distinction between connector and non-connector nodes in less segregated systems [72]. We also described the reconfiguration of redundant brain systems, finding that regions in the dorsal attention and ventral attention networks are more prone to change module assignment. Taken together, these results indicate that sensory areas form brain systems that are the most stable throughout the lifespan, but at the same time, they drive the drop of segregation of information by sharing increasingly more information with regions outside of sensory cortex.

### Technical considerations and future directions

A first technical consideration is that all the measures here reported are built upon the covariance matrices of couples of variables, so they come with an assumption of linearity of the data. Talking about redundant and synergistic interactions in this context might sound counterintuitive. However, prior works show that it is possible to search for higher-order interactions in a linear system, only capitalizing on its pairwise relationships. In [79] it has been shown that purely Gaussian systems can present higher-order synergies and that total correlation is tied to mutual information. Thus, polyadic dependencies can be observed even in linear systems. Even from [80] we know that the multivariate Gaussian is the maximum entropy distribution constrained by pairwise covariance. In other words, pairwise linear interactions set the structure upon which beyond-pairwise interactions coexist.

It is also important to note that the covariance matrices used in this paper are derived from BOLD time series of different lengths in the three datasets. In the HCP and MICA datasets we concatenated data from every subject at the node level, obtaining time series with a large number of samples; in the NKI we derived FC matrices at the subject level, from shorter time series. Information theory measures are sensitive to the length of the data on which they are computed, and from the current results it is clear that they lie in different ranges when computed on the NKI single-subject data. Thus, we pursued a simulation study to test how having fewer data points skews such measures (Figure S5). We found that TC is overestimated when computed on shorter time series and this effect worsens the larger the subset size. However, this bias is consistent across iterations such that it does not invalidate our analyses. The effect shows constant magnitude for a given length of time series, which presumably is protracted across subjects in the same way, allowing between-subject comparisons.

Another source of ambiguity is the global signal regression (GSR) performed on the BOLD time series. Previous studies suggest that global signal influences functional connectivity in a way that is spatially non-uniform, and there is a synergistic interaction between the two signals [81]. Moreover, GSR removes a large amount of redundancy present in the data [20]. Because we leveraged Gaussian assumptions to compute multivariate information theoretical measures on covariance matrices, possible confounds have to be taken into account. Nonetheless, the proposed framework is independent of the specific neural signal and could be replicated on datasets exploiting different neuroimaging modalities.

A possible limitation of our framework is that the number of communities provided as output is determined a priori by the number of communities that we impose in the seed partition. The algorithm, implemented as it is, cannot tune the spatial resolution. We tried to overcome this issue not only by examining redundant modular organization in a broad range of spatial scales, but also by attempting to identify the most interesting scales, characterized by an optimal balance between segregation and integration of information. Besides, an advantage of our framework is that it can be applied broadly to any set of multivariate time series. Its formulation is irrespective of the fact that our field of application is neuroscience and it can be tested on other complex systems, such as biological [85], ecological [86], and epidemiological [87].

This paper opens the door to a multitude of future investigations. One question is how the balance between higher-order segregation and integration is manifested in clinical cohorts. Several brain diseases are known to lack this balance [88] and multivariate information theory has already been proven relevant in clinical and behavioral contexts [89–91] Intuitively, given that neurotypical cognitive functioning requires the coordination of multiple brain regions [2, 92], beyond pairwise frameworks are needed to discover biomarkers of neurodegeneration onsets. Another line of research revolves around the relationship between structural and functional connectivity. Ebbs and flows of modular organization [93]can be reanalyzed exploiting our measures, introducing sensitivity to higher-order interactions. Finally, one can pursue investigations where our framework is modified to accommodate, for instance, a different quality function that optimizes not only the redundancy within module, but also the synergy between modules. This would require preliminary steps where a graph similar to the TSE is built for the O-information, so that it can also be compared to a chance level. Further modifications can accommodate multilayer investigations. In the case of modularity maximization, having a multilayer formulation [94] has been proven convenient to track modular structure across time [95], subjects [96], or types of connectivity [97] because nodes labeling is consistent across layers. We do believe that a similar advantage is in fact achievable with our algorithm if we optimize individual layers planting the same seed. Future investigation will elucidate the feasibility of this direction.

## METHODS

### Datasets and data processing

The analyses presented in this paper were conducted on three independent resting state fMRI datasets, derived from the Human Connectome Project (HCP) [27], the MICA57 (an open-source repository) [28], and the Nathan Kline Institute Rockland Sample (NKI) [29]. Several previous studies have used those datasets and described in detail their image acquisition and preprocessing pipelines (see for instance [98] for the HCP, [21] for MICA, and [99] for the NKI). In the following paragraphs, we recapitulate them.

#### HCP

The HCP data were derived from 100 unrelated subjects, who provided informed consent. The study protocols and procedures were approved by the Washington University Institutional Review Board. Restingstate fMRI (rs-fMRI) data were collected with a Siemens 3T Connectom Skyra (32-channel head coil) during four scans in two separate days. A gradient-echo echo-planar imaging (EPI) sequence (scan duration: 14:33 min; eyes open) was used with the following acquisition parameters: TR=720ms; TE=33.1ms; 52° flip angle; isotropic voxel resolution = 2mm; multiband factor =8.

Functional data were mapped into 200 regions covering the entire cerebral cortex, identified via the parcellation scheme proposed in [30] and aligned to the Yeo canonical resting state networks [39]. A rigorous quality control of the images was performed and led to the exclusion of 5 of the 100 unrelated subjects. Exclusion criteria included mean and mean absolute deviation of the relative root mean square (RMS) motion across resting-state fMRI and diffusion MRI scans. Four subjects exceeding 1.5 times the interquartile range of the measure distributions were excluded. One subject was then excluded because of a software error during diffusion MRI preprocessing. Downstream quality control, the data included 95 subjects, 56% female, with a mean age of 29.29 *±* 3.66.

Following a minimal preprocessing pipeline described in [100], functional images were corrected for gradient distortion, susceptibility distortion, and motion, and then aligned to the T1-weighted data. The volume was corrected for intensity bias, normalized to a mean of 10,000, then projected onto the 32*k-fs-LR* mesh, and aligned to a common space with a multimodal surface registration91. Moreover, global signal regression (GSR), detrending and band pass filtering (0.008 to 0.08HZ) were performed [101]. Finally, the first and last 50 frames of the time series were discarded, resulting in a scan length of 13.2 min and 1,100 frames.

#### MICA

The MICA dataset was approved by the Ethics Committee of the Montreal Neurological Institute and Hospital, and includes 50 unrelated subjects, who provided written informed consent. Resting-state fMRI data were collected using a 3T Siemens Magnetom Prisma-Fit (64-channel head coil). Data collections included a single scan session of 7 minutes during which participants were asked to look at a fixation cross. An EPI sequence was executed with the following acquisition parameters: TR=600ms; TE=48ms; 52° flip angle; isotropic voxel resolution = 3mm; multiband factor = 6.

Data were mapped in 200 regions of the cerebral cortex following the same parcellation scheme used for the HCP data and described in [30]. Functional images were preprocessed as outlined in [28]. Briefly, data went through motion and distortion correction, as well as FSL’s ICA FIX tool trained with an in-house classifier. Nodes were defined in the FreeSurfer surface and used to project each subject’s time series. More details about preprocessing can be found in [102], where the Micapipe93 processing pipeline is thoroughly described. Downstream, GSR was performed, as also done for the HCP data.

#### NKI

The NKI dataset consists of imaging data from a community sample of subjects encompassing a large portion of the lifespan. All participants gave written informed consent. The study received approval by the Institutional Review Boards of Nathan Kline Institute and Monclair State University. fMRI data were collected with a Siemens Magneton Trio (12-channel head coil), during a scan that lasted 9:40 seconds, where 971 participants were instructed to fixate a cross. Images were acquired using a gradient-echo planar imaging sequence with acquisition parameters set as follows: TR=645ms; TE=30ms; 60° flip angle; isotropic voxel resolution = 3mm; multiband factor = 4.

Quality control of the data included excluding subjects if the scans exceeded 1.5 interquartile range in three or more of the following metrics: DVARS standard deviation, DVARS voxel-wise standard deviation, framewise displacement mean, AFNI’s outlier ratio, and AFNI’s quality index.

Also for this dataset, the images were mapped onto 200 cerebral regions using the parcellation scheme in [30]. The fMRI data were preprocessed using the fMRIPrep version 1.1.8 [103], which comprises the briefly listed following steps, also described in its documentation. The workflow utilizes ANTs (2.1.0), FSL (5.0.9), AFNI (16.2.07), FreeSurfer (6.0.1), nipype [104], and nilearn [105]. FreeSurfer’s cortical reconstruction workflow (version 6.0) was used to skull strip the T1w, which was then aligned to the MNI space. Functional data were slice time corrected and motion corrected, using AFNI 3dTshift and FSL mcflirt, respectively. “Fieldmap-less” distortion was performed by co-registering the functional image to the same-subject T1w with intensity inverted [106] constrained with an average fieldmap template [107], implemented with antsRegistration. Then, boundary-based registration [108] was performed using bbregister to coregister the corresponding T1w. Motion correcting transformation, field distortion correcting warp, and BOLDto-T1w transformation warp were concatenated and applied altogether using antsApplyTransforms with Lanczos interpolation. For each functional run, frame-wise displacement [109] was computed using Nipype. The first four frames of the BOLD data in the T1w space were discarded. After following the fMRIPrep pipeline, images were linearly detrended, band-pass filtered (0.008-Hz), confound regressed, and standardized. Furthermore, spike regressors for frames with motion greater than 0.5mm framewise displacement were applied. Finally, GSR was performed.

### Covariance matrices definition

As in [20], the HCP-based empirical analyses presented in this paper were conducted after combining all scans and subjects to obtain a single covariance (or FC) matrix representing a grand-average of the sample. For this purpose, BOLD time series were concatenated at the node level and Pearson correlation was computed for each node pair. The same grand-average FC matrix was computed with the MICA data. With the above-chosen parcellation of the cortex, which rendered 200 nodes representing different brain regions, the average FC matrices of the HCP and MICA datasets were highly correlated (*r* = 0.851, *pval <* 10^−15^).

For the NKI-based analyses instead, we built FC matrices at the single subject level, as we wanted to be able to track individual differences linked to age. Thus, the Pearson correlation was computed for each pair of nodal time series, for each subject. After building the distribution of the mean FC values across subjects, we discarded those subjects whose mean FC exceeded the 95 percentile of the distribution. This left us with 917 covariance matrices relative to 917 subjects of age between 6 and 85 years old.

### Redundant and synergistic information in multivariate systems

In this paper, we quantified higher-order interactions by means of information theory. Specifically, given a multivariate set of signals (e.g., BOLD time series), we used the mathematics provided by information theory to measure how much the information carried by the system is shared among the variable, i.e., redundant information, and how much it banks on each variable’s contribution, i.e., synergistic information [25].

As with most measures introduced in information theory, redundancy and synergy capitalize on the basic notion of entropy [110], which quantifies the uncertainty associated with the state of a variable when only its distribution is known. Mathematically, if *X* is a discrete random variable, and *P* (*X* = *x*) is the probability distribution of its states, the entropy is formulated as follows:

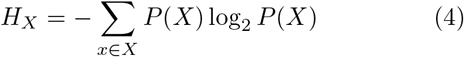

In the case of a bivariate system, where we have access to two random variables *X*_1_ and *X*_2_, entropy is used to compute the Mutual Information (I) [110], which captures how much knowing the state of the variable *X*_1_ reduces the uncertainty associated with the state of *X*_2_. Mathematically, I is formulated as follows:

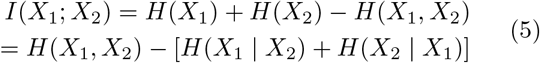

Moving towards multivariate systems then, i.e., systems comprised by more than two variables, we can quantify higher-order dependencies by extending the MI formulation [111]. There exist multiple generalization of MI. Here, we focus on three of them that will be necessary to quantify redundant and synergistic information in neural data. These are: Total Correlation (TC) [1, 33], Dual Total Correlation (DTC) [112], and Organizational Information (Ω) [24]. The TC is derived as in Eq. 6:

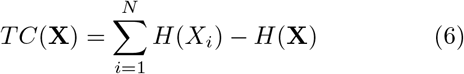

where X ={ *X*_1_, *X*_2_,…, *X*_*N*_} is a macro-variable comprising a set of *N* random variables and *H*(X) is the joint entropy of X.*TC* is low when every variable is independent and high when the joint-state of the whole system has low entropy. In other words, TC increases monotonically as the system X transitions from randomness to synchrony. TC can be used as a measure of redundancy: a multivariate system is dominated by redundant interactions when the variables share a large amount of information, hence, the state of a single variable considerably reduces the uncertainty associated with the state of every other variables, i.e., TC is high.

MI can be also generalized via the DTC as follows in Eq. 7:

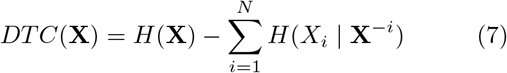

where *H*(*X*_*i*_ |*X*^−*i*^) is the residual entropy, that is the uncertainty associated with the state of the variable *X*_*i*_ that is not disclosed by the state of any other variable, or subsets of variables, comprised in **X**. With this difference, DTC captures all the entropy that is shared at least between two elements of **X**. Contrarily to TC, DTC is high in systems where *some* information is shared, but is low in both cases where **X** comprises totally random or synchronized variables.

The organizational information is the difference between TC and DTC (Eq. 8), so that when it is positive (TC(**X**)>DTC(**X**)) redundant information among the variables is predominant, whereas when it is negative (TC(**X**)<DTC(**X**)) the system is characterized by information that is both shared but not redundant.

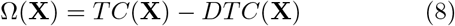

Ω is low in systems dominated by synergistic interactions and high in systems dominated by redundant interactions, and we used it as a proxy for synergy in our empirical analyses. For a thorough discussion on how the O-information can be interpreted see also [113, 114].

### Information theory in Gaussian systems

All the information theoretic measures reported above require the estimation of the entropy H(X), and specifically of P(x), which can be challenging on empirical continuous data, like fMRI BOLD signals. However, for normally distributed (Gaussian) multivariate systems, closed-form estimators exist [115] that make the computation of the joint entropy easier. Under the assumption that BOLD data follows a multivariate Gaussian distribution – an assumption supported by numerous studies [116–118] – we can exploit those close-form estimators and derive information theoretical measures directly from the covariance matrix.

For a single Gaussian random variable *X* ∼ *N* (*µ, σ*), the entropy can be computed as:

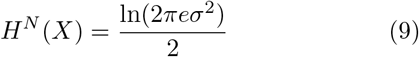

For a multivariate Gaussian random variable **X** ={*X*_1_, *X*_2_,…, *X*_*N*_}, the joint entropy can be computed as:

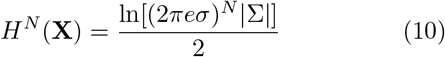

where| Σ | denotes the determinant of the covariance matrix of **X**.

Similarly to what has been shown in the previous section, we can generalize these formulations of entropy and derive the MI of a bivariate system comprising the variables *X*_1_ and *X*_2_. Then, given the Pearson correlation coefficient between *X*_1_ and *X*_2_, here referred to as *ρ*, the MI can be written as:

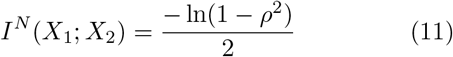

Finally, the estimator for the total correlation of a multivariate Gaussian system has the following formulation:

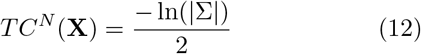

Then, from these equations, it is straightforward to rewrite in close-form both the dual total correlation and the organizational information. For a more thorough description of how information theoretical measures can be drawn in close-form see also [20, 115].

### Multiscale modularity maximization on brain etworks

Modular structure was inferred on the HCP dataset via modularity maximization [31], which, given an adjacency matrix, returns partitions of the networks into modules that are maximally internally dense with respect to a chance level (or null model). Modularity is commonly indicated with Q and is formulated as follows:

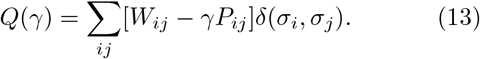

where *W*_*ij*_ and *P*_*ij*_ are the weights of the connections between node *i* and node *j* in the adjacency matrix *W* and in the null model *P, γ* is a resolution parameter, and *δ*(*σ*_*i*_, *σ*_*j*_) is a factor whose value is 1 when *i* and node *j* belong to the same community and 0 otherwise.

The choice of the null model P is non-trivial. The weight that we expect to find between two nodes strongly depends on the characteristics of the adjacency matrix *W* [119]. For covariance matrices, i.e., full weighted matrices, a reasonable choice is the Potts null model, where *P*_*ij*_ holds the same value for every pair of nodes and is modulated by *γ*. The choice of *γ* then, defines the spatial resolution of the partition in terms of size and number of modules, so that low *γ*-values yield to a coarse modular structure, whilst high *γ*-values produce a finer modular structure.

In this paper, we applied a 2-step multiresolution approach, called multiresolution consensus clustering (MRCC) [37]. In the first step, we linearly sampled 1000 values of *γ* in the range [0, 1] and we ran modularity maximization with each one of these *γ*-values obtaining 1000 differently resolved partitions. At this point, two *γ*-values have been identified, *γ*_*L*_ and *γ*_*H*_, which generated partitions with a number of modules between 2 and *N/*4 (with *N* = 200 being the number of nodes). Then, in the second step, we linearly sampled 1000 values of *γ*, but this time in the range [*γ*_*L*_, *γ*_*H*_], running modularity maximization with each of these values. Again, retaining partitions with at least 2 and maximum *N/*4 modules, we obtained 990 partitions (10 *γ*-values/partitions did not survive the limits imposed on clusters number) at different spatial resolutions that we used for our empirical analyses.

## DATA AND CODE AVAILABILITY

Network data and MATLAB code to reproduce the analysis are available at https://github.com/mariagraziaP/comm_detection_via_redundancy_optimization/tree/main.

## ACKNOWLEDGMENTS

This research was supported in part by Lilly Endowment, Inc., through its support for the Indiana University Pervasive Technology Institute which provided the computer infrastructure required for completion of the project. The funders had no role in study design, data collection and analysis, decision to publish, or preparation of the manuscript.

---

**FIG. S1.**
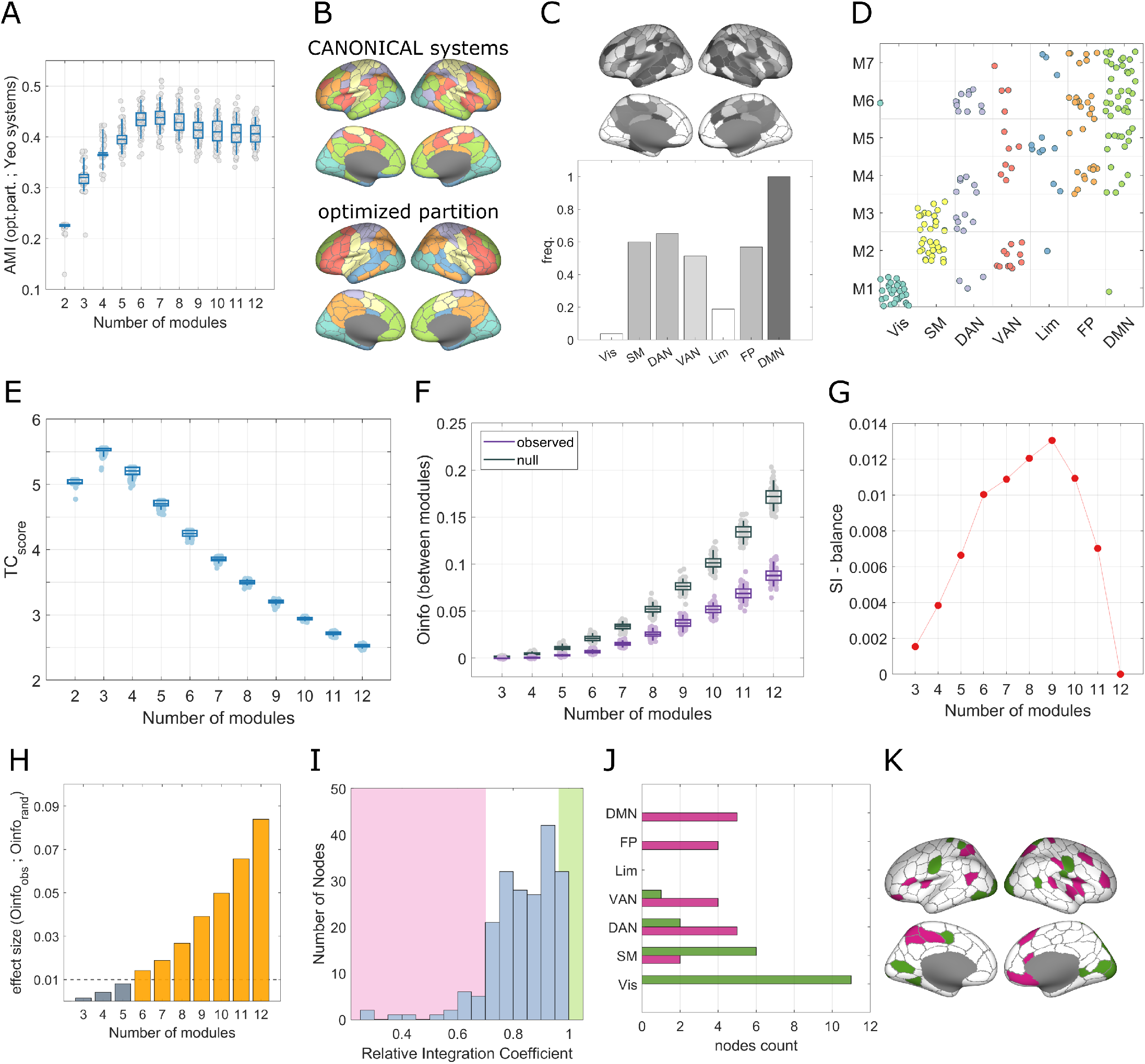
Replication of the analyses on the MICA dataset. **A**. Distance between the canonical systems and the partitions obtained optimizing TC_score_ on the MICA dataset, in terms of adjusted mutual information (AMI). A peak has been found in partitions with 7 modules, where AMI is on average maximum. **B**. Projection on the cortex surface of the canonical systems (up) and of an example of redundancy dominated partition with 7 modules (down). **C**. Frequency with which nodes change module allegiance between the 7-modules partitions and the canonical system across 100 iterations of the TC_score_ maximization. **D**. Mapping of each node module allegiance in the partitions shown in panel B. **E-H**. Trends across spatial resolution of TC_score_, O-information, Effect Size (between the observed and the null O-information), and B. **I**. Distribution of RIC values computed for every node of the 7-modules centroid partition. **J-K**. Mapping of segregator (green) and integrator (pink) nodes among the canonical systems and on the cortex surface.

**FIG. S2.**
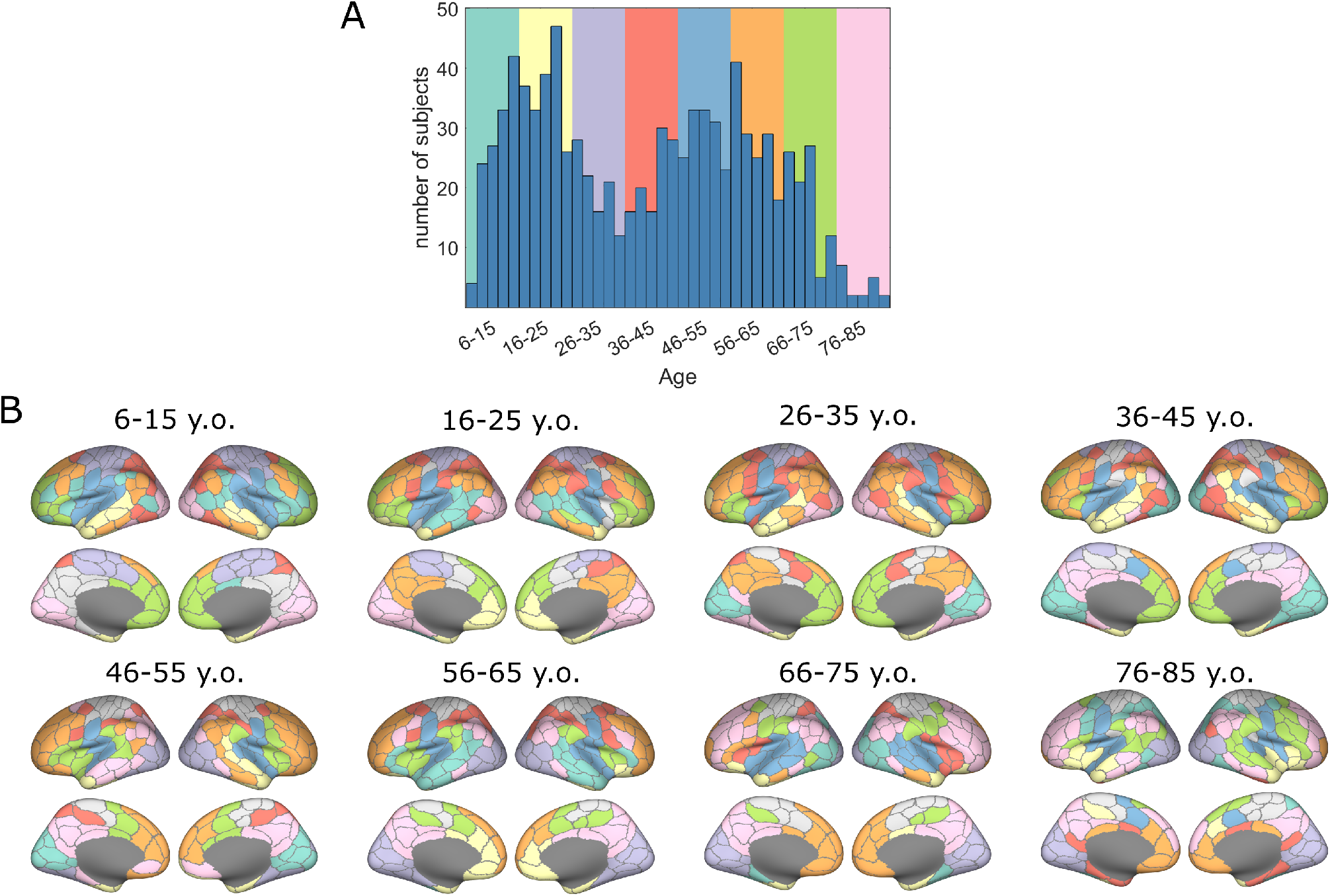
**A**. Distribution of the NKI population across the lifespan. The colored rectangles in the background define the age-groups identified for the second half of the analyses presented in the main text. **B**. Mapping on the cortex of the centroid partitions derived from each age group.

**FIG. S3.**
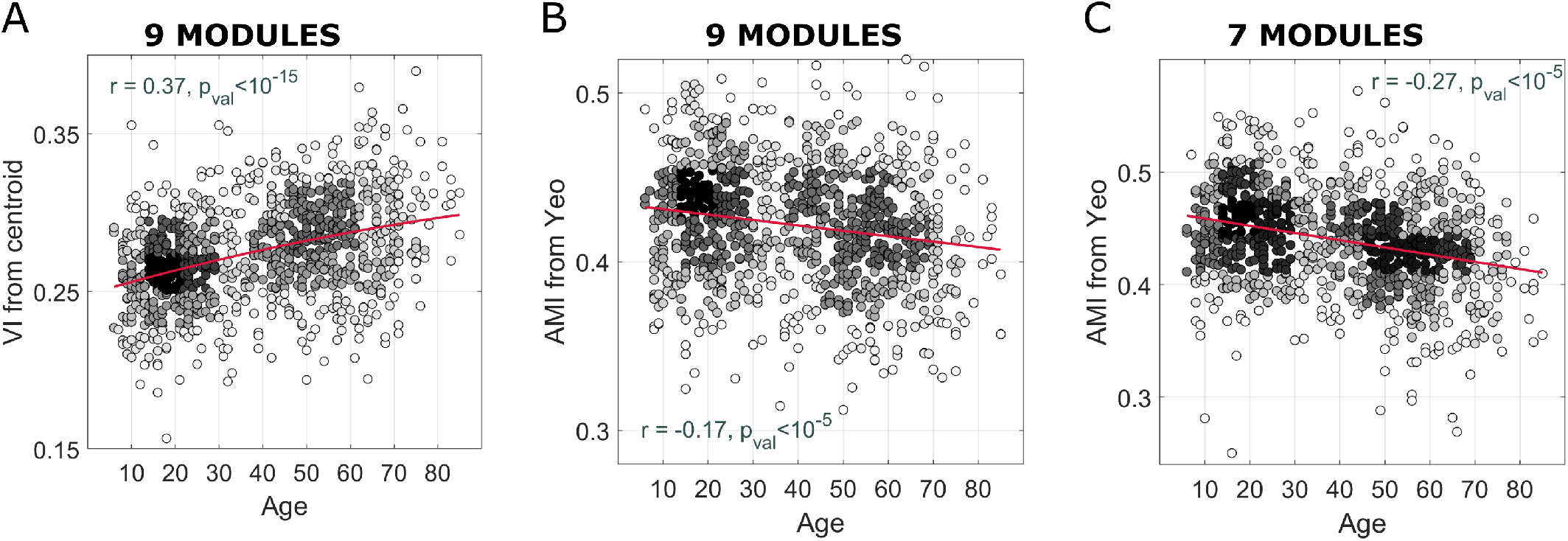
**A**. Having 100 iterations of the TC_score_ maximization for each subject, we plot the average variation of information (VI) between the centroid partition and all the other 99 partitions for each subject. Because there is a positive correlation between this VI and age, we can conclude that aging is associated with less stability of the algorithm. **B-C**. Distance, in terms of adjusted mutual information (AMI), between the canonical systems and the TC_score_ optimized partitions across the lifespan. The negative correlation denote an increasing divergence from the canonical system associated with age.

**FIG. S4.**
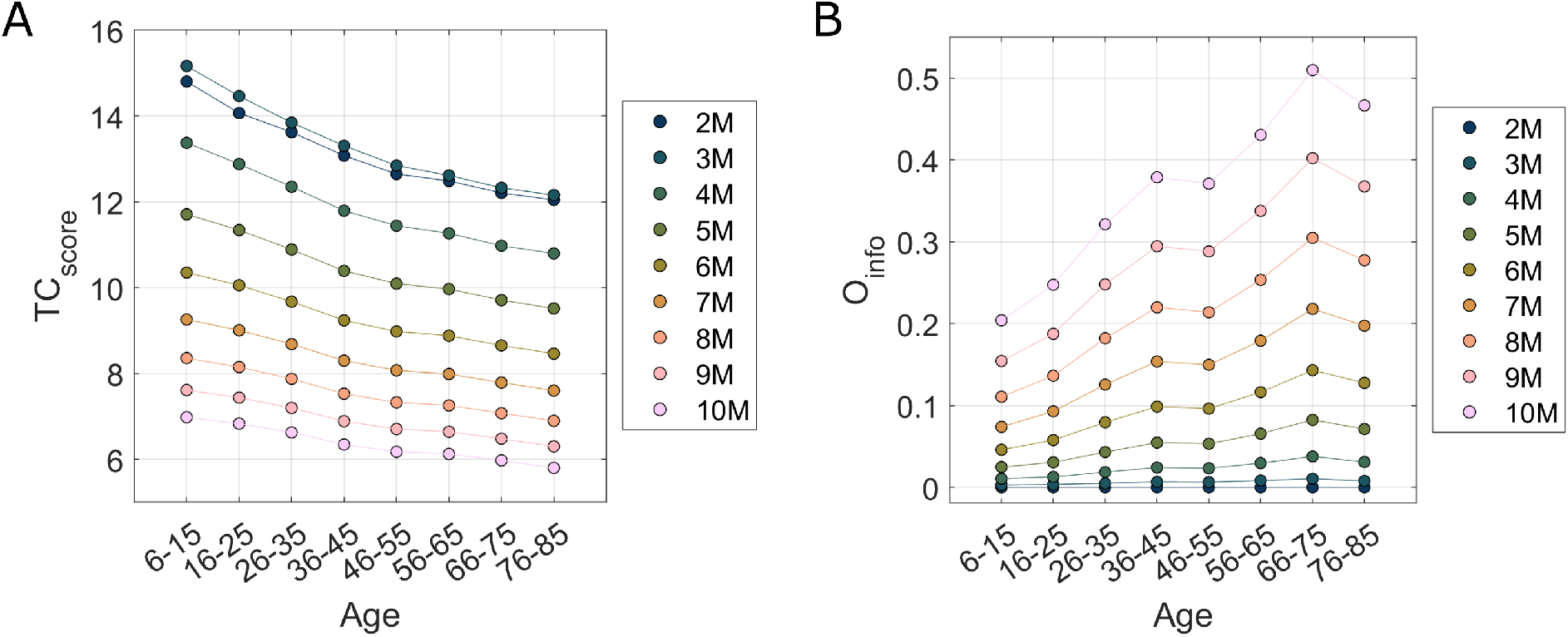
Trends across the lifespan of the TC_score_ and O-information for partitions with different spatial resolution, presenting from 2 to 10 modules. In order to have a clearer visualization, we reported average values computed resampling TC_score_ and O-information within age-groups.

**FIG. S5.**
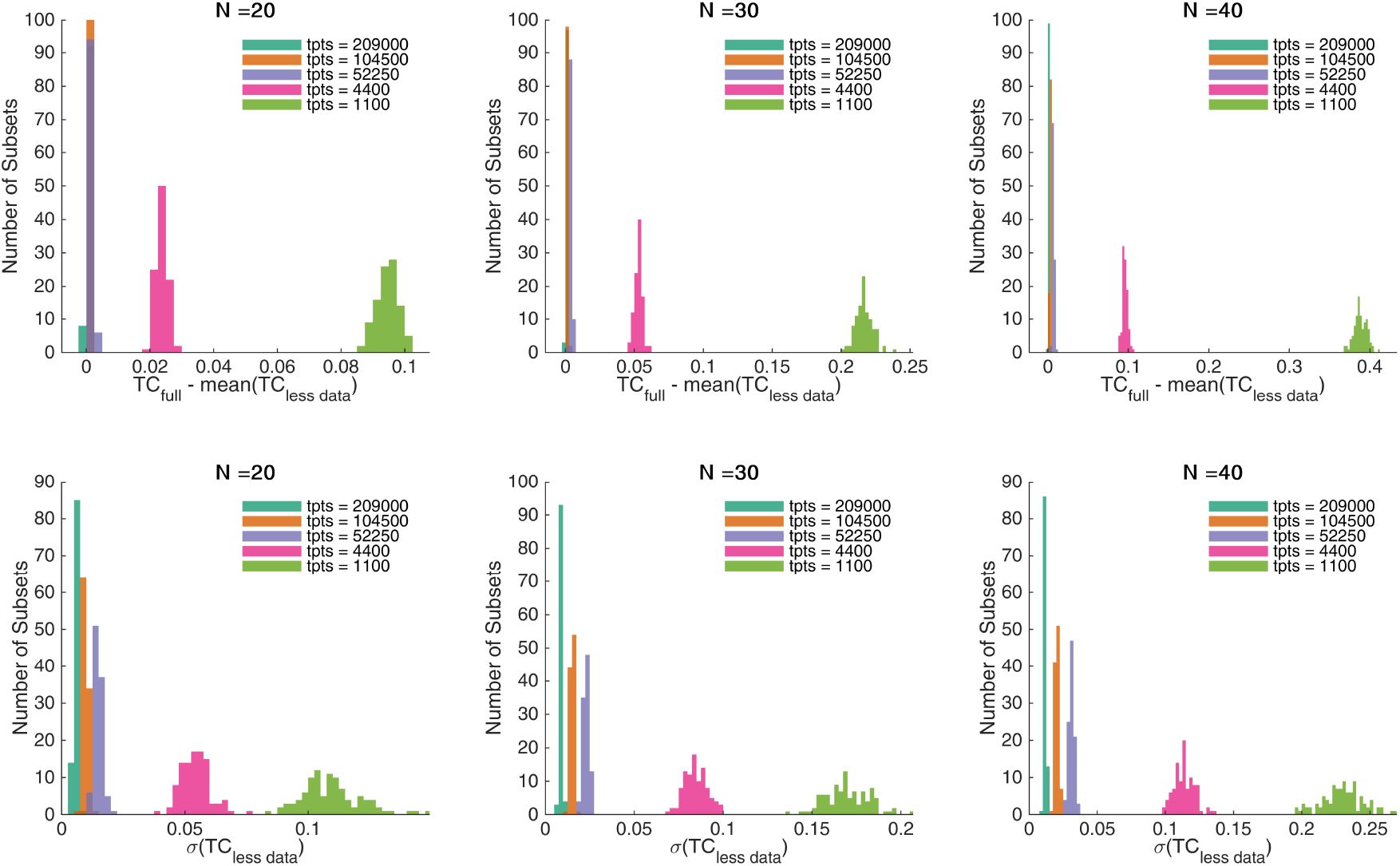
Starting from the HCP combined time series, we tested subsets of different sizes represented by columns (N=20, 30, and 40) and for each one of them we recalculated the TC keeping progressively fewer data points: half of the data points (blue), then only a quarter (orange), then an eight (yellow), then 4400 (purple), and then 1100 (green), which is the length of a single scan. The random data points were dropped 1000 times per subset. In the first row, we report the difference between the TC computed on the full dataset and the average TC computed on each subset’s 1000 randomizations with less data. The more time points are discarded, the more TC results overestimated. Moreover, the overestimation becomes worse with increasing subset size. In the second row, we report the standard deviation of the TC computed on different fractions of the data for each subset. TC values vary more when less data is included in the calculation. Again, this phenomenon worsens with increasing subset size.

